# Widespread heteroresistance to antibiotics in Lactobacillus species

**DOI:** 10.1101/2025.03.24.644958

**Authors:** Dormarie E. Rivera-Rodriguez, Chayse Busby, Luisa Cervantes-Barragan, David S. Weiss

## Abstract

Lactobacilli are prevalent members of the intestinal and reproductive tract microbiota of humans and other species. They are commonly used in probiotics and various food products due to their beneficial effects on human health. For example, these beneficial microbes are used to treat diarrhea caused by antibiotic therapy and are commonly given during antibiotic treatment. Despite the many studies conducted to understand the beneficial effects of Lactobacilli, less is known about their resistance and heteroresistance to antibiotics. In this study, we evaluated the resistance heterogeneity in eight *Lactobacillus* species. Our results demonstrate that several Lactobacilli species, including *Lactobacillus rhamnosus*, are heteroresistant to antibiotics, a recently discovered phenotype commonly seen in multidrug-resistant organisms that cause clinical failures but understudied in commensals and probiotics.

## Introduction

The intestinal microbiota is composed of ∼38 trillion microorganisms. Several members of the microbiota confer intestinal protection, for example, by adhering to the mucous layer and preventing pathogens from colonizing the gut or by activating protective immune responses[1]. Some also metabolize dietary products for the host to use as an energy source. For example, members of the *Lactobacillaceae* family, a major family residing in the ileum of the small intestine and other mucosal sites, express mucus-binding proteins that help them selectively adhere to the intestinal mucosa[2-4]. These facultative anaerobes can also help metabolize sugars, such as lactose, and other dietary components[1, 2, 5-10].

Due to their benefits to human health, commensal microbes, have been used as probiotics. Lactobacillus species are some of the most widely used probiotics, and are well known for their beneficial effects, including treating antibiotic-associated diarrhea and intestinal inflammatory diseases[9, 11], and are often used during antibiotic treatment. Previous studies have shown that Lactobacilli can have preexisting resistance to various antibiotics, possibly due to intrinsic resistance mechanisms or acquired resistance[8, 12-22]. These studies have assessed the genomic and phenotypic resistance mechanisms in Lactobacilli. However, if Lactobacilli can also exhibit heteroresistance to various antibiotics remains unknown.

Here, we systematically analyzed the antibiotic resistance of eight Lactobacillus species to a selection of antibiotics from the most common antibiotic classes: beta-lactams, tetracyclines, glycopeptides, fluoroquinolones, macrolides, and aminoglycosides. Our results show that while some Lactobacillus species are resistant to antibiotics, several are also heteroresistant, a recently discovered type of resistance commonly seen in multidrug-resistant organisms that cause clinical failures but are understudied in commensals and probiotics. Interestingly, *L. rhamnosus*, a commonly used in probiotic, exhibits heteroresistance against fluoroquinolones and some cephalosporins. We further tested the *L. rhamnosus* heteroresistant phenotype to ciprofloxacin by performing an antibiotic resistance stability assay, which confirmed *L. rhamnosus* was indeed heteroresistant to ciprofloxacin. Our study provides a comprehensive analysis of antibiotic resistance to antibiotics in the Lactobacillus genera and describes the presence of heteroresistant Lactobacillus species.

## Materials and Methods

### *Lactobacillus* isolates and culture conditions

*Lactobacillus acidophilus* 4356, *L. plantarum* 9014, *L. gasseri* BAA-2841, *L. helveticus* 15009, and *L. murinus* 35020 were purchased from the American Type Culture Collection (ATCC). *L. johnsonii* WU was isolated from mice housed at Washington University. *L. reuteri* 100-23 (ID: 17509) strain was obtained from the German Collection of Microorganisms and Cell Cultures (DSMZ). *L. rhamnosus* was isolated from probiotics tablets. Lactobacilli from frozen glycerol stocks were struck onto De Mann Rogosa and Sharpe (MRS) (BD Difco, Cat. No. 288210) agar plates. Single colonies were inoculated into MRS broth (BD Difco, Cat. No. 288130) with Tween80 (Sigma, Cat. No. P8074-500mL). Cells were incubated overnight (12-16 hours) at 37°C under aerobic or anaerobic conditions.

### Population Analysis Profile assays

Population analysis profiles (PAPs) were conducted as previously described[23-25]. Solid agar plates were made for each antibiotic at seven concentrations using MRS agar. Concentrations were calculated to contain 0.06, 0.12, 0.25, 0.50, 1.00, 2.00, and 4.00 times the breakpoint for each antibiotic **(Table S1)**. As reference we used the breakpoint concentrations for *Enterococcus sp*. from the Clinical & Laboratory Standards Institute (CLSI) Guidelines: Ciprofloxacin 4µg/mL; Levofloxacin 8µg/mL; Delafloxacin 4µg/mL; Ampicillin 16µg/mL; Amoxicillin 16µg/mL; Penicillin 16µg/mL; Erythromycin 8µg/mL; Azithromycin 8µg/mL; Tetracycline 16µg/mL; Minocycline 16µg/mL; Vancomycin 32µg/mL. We used these breakpoints because they resemble the relative minimal inhibitory concentration (MIC) used to treat *Enterococcus* infections[26-29]. If no concentrations were found for Enterococcus sp. in CLSI, we used the breakpoint concentrations of pathogenic Anaerobes from CLSI: Cefepime 16µg/mL; Ceftriaxone 64µg/mL; Cefazolin 64µg/mL. We used these breakpoints from pathogenic anaerobes, such as *Clostridium difficile*, due to similar MIC concentrations used to treat *C. difficile* infections[29-31]. No concentrations were found for *Enterococcus sp*. and *C. difficile* for aminoglycoside concentrations. Thus, we used the following concentrations from previous studies[32]: Streptomycin 16µg/mL, Gentamycin 16µg/mL, and Neomycin 16µg/mL. Single colonies from Lactobacilli frozen stocks were inoculated and grown overnight in MRS broth. Each Lactobacilli isolate was serially diluted the following day using 1X phosphate-buffered saline (PBS). Serial micro-dilutions were plated in antibiotic-free plates or each antibiotic concentration plate. Colonies were enumerated after a 48-hour incubation period at 37°C under aerobic or anaerobic conditions. An isolate was classified as resistant (R) if the colony-forming units (CFU) that grew at the breakpoint concentration were at least 50% of those that grew on the antibiotic-free plates. If an isolate was not R, they were classified heteroresistant (HR) if the CFUs that grew at 2 or 4 times the breakpoint was at least 0.0001% (1 in 10^6^) of those that grew in the antibiotic-free plates. If the isolate was not R or HR, they were classified as susceptible (S). (**Figure S1A**)

### Antibiotic resistance stability assay

Antibiotic resistance stability assay was conducted as previously described[23]. The following concentrations were used: Ciprofloxacin (4µg/ml) and Levofloxacin (8µg/ml). Briefly, *L. murinus* or *L. rhamnosus*, categorized as HR to ciprofloxacin and levofloxacin respectively, were cultured in MRS media at 37°C under aerobic conditions for 24 hours. Afterwards, they were sequentially subcultured (1:100) in the presence of ciprofloxacin or levofloxacin at 37°C under aerobic conditions every 24hr for 48 hours. After subculturing on antibiotic containing media for 48 hrs, antibiotic-free subcultures were performed by diluting antibiotic-treated cultures (1:1000) and incubating for 24 hours at 37°C under aerobic conditions. These antibiotic-free subcultures were continued until fourteen generations of growth without drugs. Cells were serially diluted in 1X PBS and plated on MRS agar with or without the respective antibiotic to determine the proportion of resistant bacteria after each subculture.

### Statistical analysis

All experiments were repeated with biological triplicates at least thrice to ensure reproducibility. Error bars represent the standard error of the mean (SEM) from biological triplicates. Statistical analysis was performed using GraphPad Prism software.

## Results

### Lactobacilli exhibit resistance heterogeneity against different antibiotic classes

While previous studies have provided insights into the resistance phenotypes of various Lactobacillus species[11-13, 20, 21, 32-34], the specific question of whether Lactobacilli exhibit varied resistance to different antibiotic classes, including heteroresistance, remains largely unexplored. We systematically analyzed the resistance to antibiotics in the Lactobacillus genera. To this end, we selected eight Lactobacillus species known to be members of the human and mouse microbiome, used as probiotics, or present in fermented foods. Using the population analysis profile (PAP) method, we tested their resistance profile to antibiotics of the most commonly used antibiotic classes; four bactericidal (beta-lactams, aminoglycosides, glycopeptides, and fluoroquinolones) and two bacteriostatic (tetracyclines and macrolides).The PAP method quantifies the proportion of bacterial growth on seven increasing concentrations of the tested antibiotic and selected to contain 0.06, 0.12, 0.25, 0.50, 1.00, 2.00, and 4.00 times the breakpoint for the antibiotic. The PAP curves for each isolate and antibiotic were classified as resistant, heteroresistant, or susceptible based on predetermined criteria and as described in the Materials and Methods and the supplemental figure (**Figure S1A**).

### Resistance phenotype in bactericidal antibiotics

#### Beta-lactams

Studies using antibiotic susceptibility assays showed that Lactobacilli are susceptible to different beta-lactams, mainly penicillins[12, 13, 22, 34, 35]. Thus, we first focused on characterizing resistance levels to beta-lactams, such as penicillins and cephalosporins. Using the PAP method. Consistent with previous reports, all tested Lactobacilli were susceptible to amoxicillin (**Figure 1B**), penicillin (**Figure S1D**), and ampicillin (**Figure S1E**). Interestingly, our data demonstrates that *L. murinus* is resistant to cefepime (**Figure 1C**), heteroresistant to ceftriaxone (**Figure S1F**), and susceptible to cefazolin (**Figure S1G**). Additionally, other Lactobacillus species, such as *L. reuteri* and *L. acidophilus*, were heteroresistant to cefepime (**Figure 1C**) but were susceptible to ceftriaxone (**Figure S1F**) and cefazolin (**Figure S1G**). These results suggest that Lactobacilli can be heteroresistant to certain beta-lactams, such as cephalosporins.

**Figure 1:**
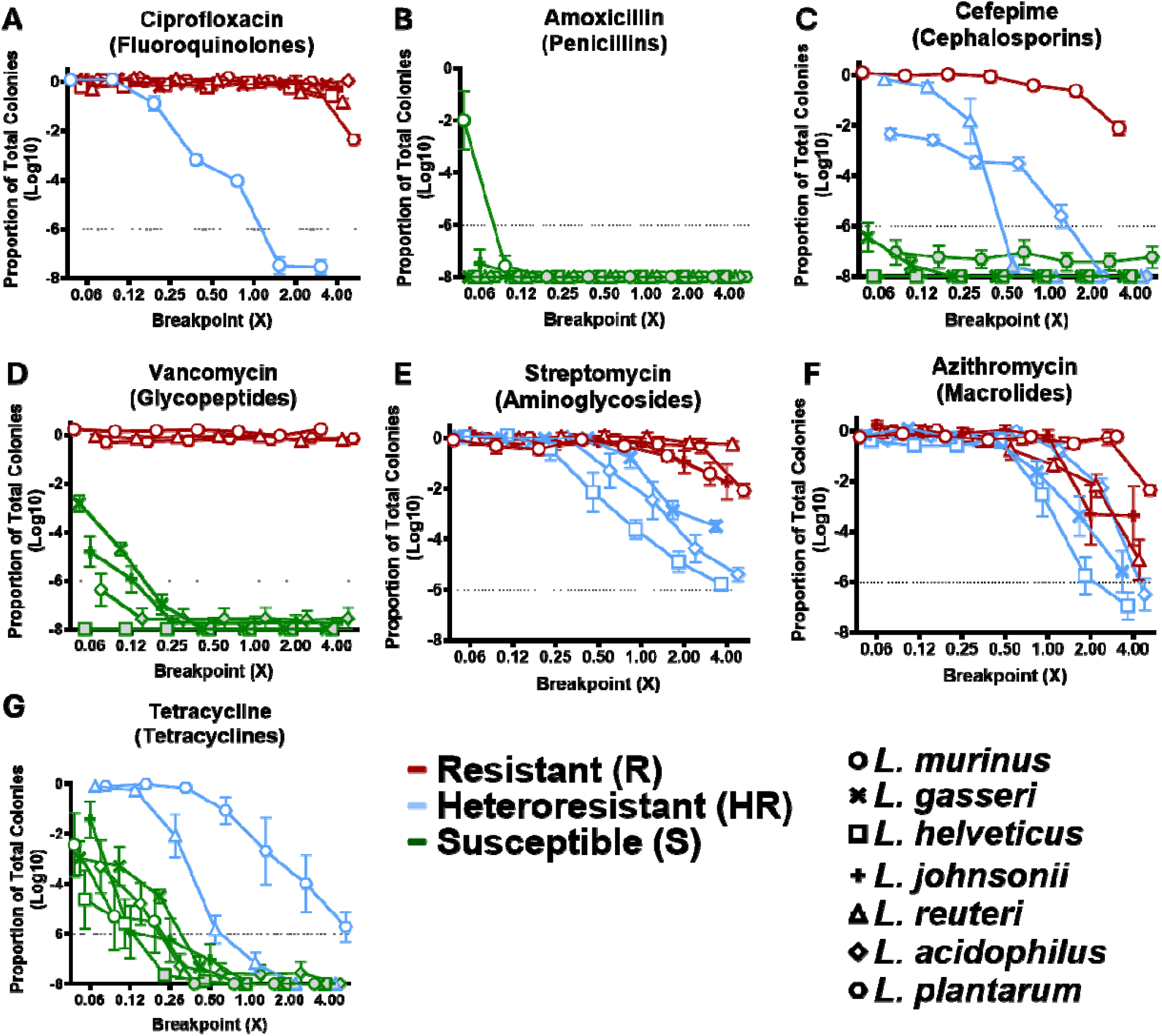
Lactobacilli have varied resistance to different antibiotics. Population analysis profile assays (PAPs) of seven Lactobacillus isolates plated on **A)** Ciprofloxacin (MIC: 4µg/ml), **B)** Amoxicillin (MIC: 16µg/ml), **C)** Cefepime (MIC: 16µg/ml), **D)** Vancomycin (MIC: 32µg/ml), **E)** Streptomycin (MIC: 16µg/ml), **F)** Azithromycin (MIC: 8µg/ml), **G)** Tetracycline (MIC: 16µg/ml) in aerobic conditions. **Gray-filled** dots indicate samples below the level of detection. Symbols represent bacterial species. Graphs are representative of 3 independent experiments (n=3).

#### Fluoroquinolones

Our data showed that six out of eight Lactobacilli were resistant to ciprofloxacin, a second-generation fluoroquinolone[36] (**Figure 1A**). However, *L. murinus* was heteroresistant to ciprofloxacin in aerobic (**Figure 1A**) and anaerobic conditions (**Figure S2A & S2B**). Since most Lactobacilli were resistant to ciprofloxacin, we investigated if Lactobacilli were also resistant to newer fluoroquinolones, such as levofloxacin, a third-generation fluoroquinolone, and delafloxacin, a fourth-generation fluoroquinolone[36]. Interestingly, only *L. gasseri, L. helveticus*, and *L. acidophilus* were resistant to levofloxacin, whereas the remaining Lactobacilli were heteroresistant (**Figure S1B**). Moreover, all Lactobacilli were susceptible to delafloxacin (**Figure S1C**). We conclude that most Lactobacilli are heteroresistant to ciprofloxacin and levofloxacin; however, they are susceptible to newer fluoroquinolones, such as delafloxacin.

#### Glycopeptides

Studies from Zhang and others showed that most Lactobacillus isolates encode vanX. This gene encodes the Ddl enzyme, which alters the peptidoglycan residues in their D-alanine terminus to prevent vancomycin binding, making them intrinsically resistant to vancomycin[22, 34, 37, 38]. Using the PAP method, our results confirmed that *L. murinus, L. reuteri*, and *L. plantarum* were resistant to vancomycin, further validating previous studies[22, 34, 35, 38]. However, *L. gasseri, L. helveticus, L. johnsonii*, and *L. acidophilus* were susceptible to vancomycin (**Figure 1D**). These results correlate with previous reports showing that *L. gasseri, L. johnsonii*, and *L. acidophilus* are more sensitive to vancomycin[12, 13, 37].

#### Aminoglycosides

It has been previously reported that Lactobacilli are also intrinsically resistant to aminoglycosides due to the lack of cytochrome-mediated drug transport[13, 32, 34]. Thus, we expected all Lactobacilli used in this study to resist different aminoglycosides. To our surprise, three *Lactobacillus* species, *L. gasseri, L. helveticus*, and *L. acidophilus*, were heteroresistant to streptomycin, while the remaining Lactobacilli were resistant (**Figure 1E**). Only *L. helveticus* and *L. plantarum* were heteroresistant to gentamycin, while the others were resistant (**Figure S1H**). Nevertheless, all eight Lactobacillus species resisted neomycin (**Figure S1I**), confirming previous research using antibiotic susceptibility assays[12, 13].

### Resistance phenotype in bacteriostatic antibiotics

#### Tetracyclines

Previous studies show that Lactobacilli are susceptible to tetracyclines[12, 13, 22, 33]. However, these studies commonly use antibiotic susceptibility assays to determine resistance. Using the PAP method, we observed most of the Lactobacillus species are susceptible to tetracycline (**Figure 1G**) and minocycline (**Figure S1K**). *L. plantarum* was heteroresistant to both antibiotics (**Figure 1G & S1K**), while *L. reuteri* is only heteroresistant to tetracycline (**Figure 1G and S2**).

#### Macrolides

Previous studies suggest some Lactobacilli to be susceptible to macrolides since they are generally susceptible to inhibitors of protein synthesis[13, 20, 35]. However, many others reported that Lactobacilli can have acquired resistant mechanisms against macrolides[13, 22, 33, 39, 40]. Thus, we used our PAP method to determine the macrolide resistance phenotype in Lactobacilli. Surprisingly, *L. gasseri, L. helveticus*, and *L. acidophilus* were heteroresistant to azithromycin (**Figure 1F**), whereas all eight Lactobacilli were resistant to erythromycin (**Figure S1J**).

### Lactobacilli species can be heteroresistant to antibiotics

To confirm that Lactobacilli are truly heteroresistant and not persister cells, which do not robustly replicate in antibiotic-treated media[23, 41], we selected two bacteria-antibiotic pairs that exhibit an heteroresistant phenotype and performed an antibiotic resistance stability assay. In this assay, bacteria which are heteroresistant to an antibiotic will increase in their proportion of resistant cells when subcultured in the presence of antibiotic, but the proportion of resistant cells will decrease again when further subcultured in the absence of antibiotic, in contrast to persisters, or cells with newly acquired resistance which will remain high. Our results demonstrate that the resistant subpopulation in *L. murinus* rapidly increased in ciprofloxacin containing cultures. However, when further subcultured without ciprofloxacin, the frequency of resistant subpopulation decreased after two weeks (**Figure 2**). A similar result was obtained when testing *L. plantarum* heteroresistance to Levofloxacin (**Figure S3**). These results confirm that Lactobacillus species can be heteroresistant to antibiotics.

**Figure 2:**
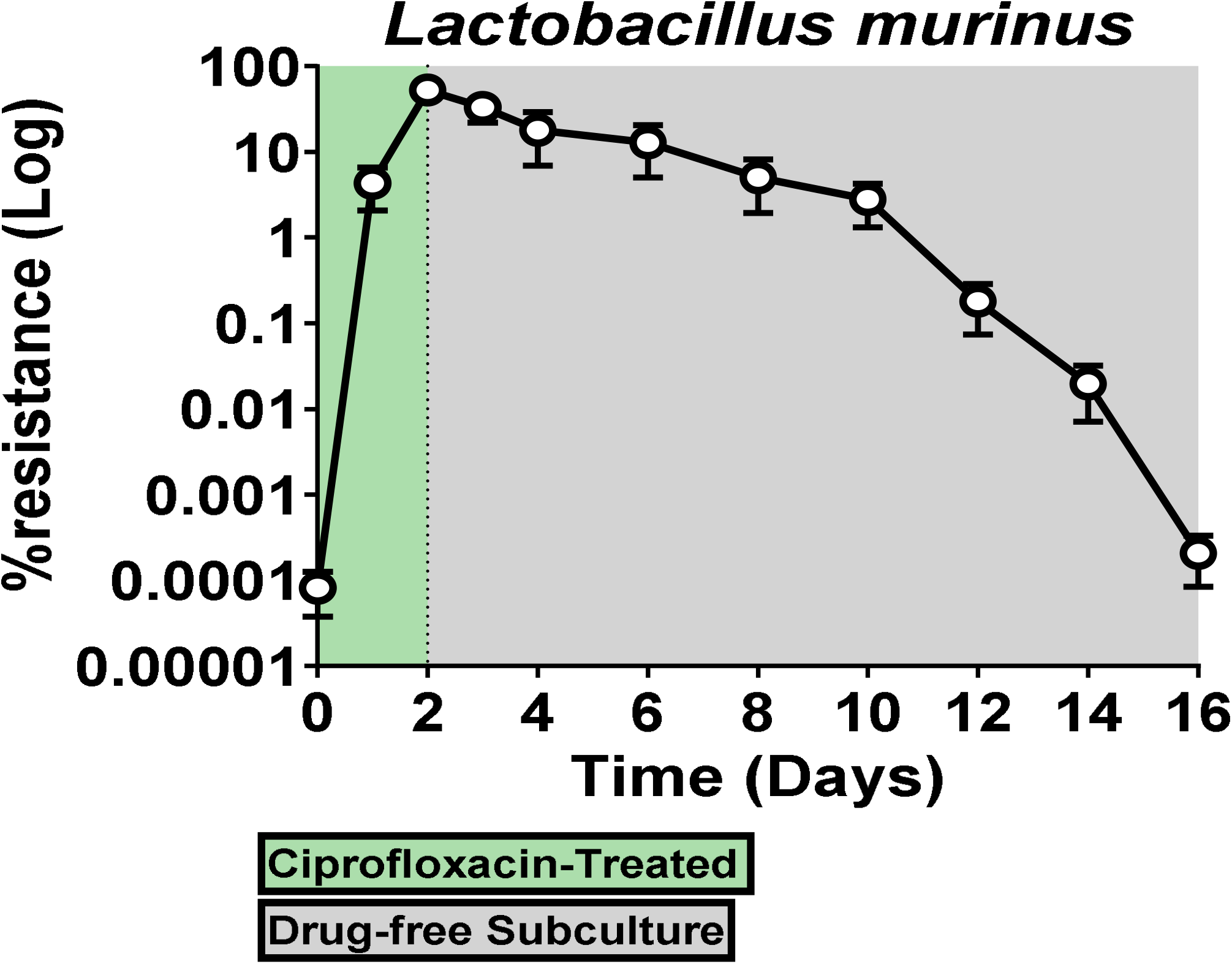
*Lactobacillus murinus* is heteroresistant to ciprofloxacin. *Lactobacillus murinus* was subcultured in MRS with ciprofloxacin (MIC: 4µg/ml) for 48 hours followed by subcultures in ciprofloxacin-free MRS media for 14 days. Data represents the proportion of ciprofloxacin resistant bacteria in the culture each day. Symbols represent the mean of triplicates and is representative of three independent experiments (n=9).

### *L. rhamnosus* is heteroresistant to Fluoroquinolones Ciprofloxacin and Levofloxacin

Probiotics are live bacteria safe for consumption and used to improve health[22]. Previous studies have reported that many Lactobacilli found in probiotics, such as *L. rhamnosus*, can resist various antibiotics[13, 22]. We decided to test *L. rhamnosus* antibiotic resistance and determine if it was also heteroresistant to antibiotics. Using the PAP method on a *L. rhamnosus* we determined it was susceptible to all tested penicillins (**Figure 3B, S4C & S4D**), to the cephalosporins, cefazolin (**Figure S4E)** and ceftriaxone (**Figure S4F**), to delafloxacin (**Figure S4B**), and the tested tetracyclines (**Figure 3G & S4J**). *L. rhamnosus* was resistant to cefepime (**Figure 3C**), vancomycin (**Figure 3D**), and all tested aminoglycosides (**Figure 3E, S4G & S4H**), and macrolides (**Figure 3F & S4I**). Interestingly, *L. rhamnosus* was heteroresistant to the fluoroquinolones ciprofloxacin (**Figure 3A**) and levofloxacin (**Figure S4A**). We confirmed the heteroresistant *L. rhamnosus* to ciprofloxacin (**Figure 3H**) or levofloxacin (**Figure S4K**) using the antibiotic resistance stability assay. Our results demonstrate that the proportion of resistant subpopulations in *L. rhamnosus* rapidly increased in ciprofloxacin (**Figure 3H**) and levofloxacin (**Figure S4K**). Subsequently, when subcultured without antibiotics, the frequency of each resistant subpopulation decreased after two weeks (**Figures 3H & S4K**) confirming that *L. rhamnosus* is heteroresistant to ciprofloxacin and levofloxacin. Overall, our results provide a comprehensive analysis of the antibiotic resistance and heteroresistance of several Lactobacillus species to antibiotics from all major classes of antibiotics and demonstrate that commensal bacteria and probiotics such as *L. rhamnosus* can also exhibit heteroresistance.

**Figure 3:**
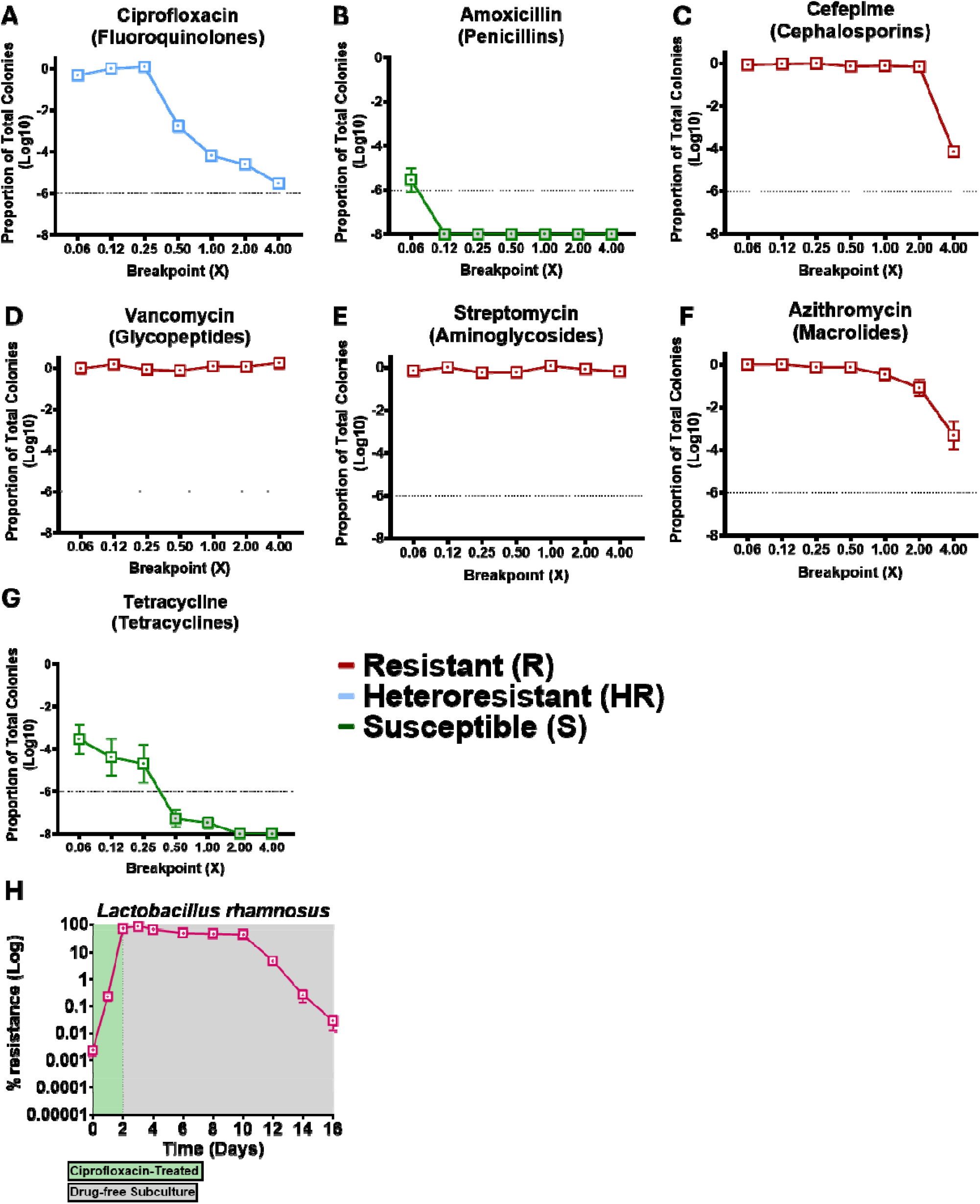
*L. rhamnosus* is heteroresistant to Fluoroquinolones Ciprofloxacin and Levofloxacin. Population analysis profiles assay (PAPs) of *Lactobacillus rhamnosus* plated on **A)** Ciprofloxacin (MIC: 4µg/ml), **B)** Amoxicillin (MIC: 16µg/ml), **C)** Cefepime (MIC: 16µg/ml), **D)** Vancomycin (MIC: 32µg/ml), **E)** Streptomycin (MIC: 16µg/ml), **F)** Azithromycin (MIC: 8µg/ml), **G)** Tetracycline (MIC: 16µg/ml) in aerobic conditions. Gray-filled dots indicate samples below the level of detection. PAPs were performed in triplicates Graphs are represewntative of 3 independent experiments. **H)** *L. rhamnosus* was subcultured in MRS with ciprofloxacin (MIC: 4µg/ml) for 48 hours followed by subcultures in ciprofloxacin-free MRS media for 14 days. Data represents the proportion of ciprofloxacin resistant bacteria in the culture each day. Symbols represent the mean of triplicates and are representative of three independent experiments (n=9).

## Discussion

*Lactobacillus* species are beneficial microbes that help the host metabolize dietary products and prevent pathogen colonization by selectively adhering to the intestinal mucosa[1-8, 42, 43]. These bacteria are commercially used in various dairy and meat products and are some of the most commonly used bacteria in probiotic supplements[9, 10]. Probiotics containing Lactobacilli are commonly prescribed to patients with antibiotic-associated diarrhea and dysbiosis[9, 11] and are commonly consumed during or shortly after antibiotic therapy. Previous studies have analyzed the resistance or susceptibility of Lactobacillus species to a wide array of antibiotics. However, little is known about the presence of heteroresistance in commensals and probiotics, a recently discovered phenotype commonly seen in multidrug-resistant organisms.

Along with previous studies[12, 13, 22, 34, 35], we confirmed that Lactobacilli are susceptible to penicillin, ampicillin, and amoxicillin. Interestingly, many Lactobacilli exhibit heteroresistance to various cephalosporins. For example, *L. reuteri* and *L. acidophilus* were heteroresistant to cefepime, whereas *L. murinus* was heteroresistant to ceftriaxone. Recent work on carbapenem-resistant *Lactobacillus* isolates from clinical case reports and starter cultures showed that cephalosporin resistance levels are highly variable in Lactobacilli[12, 22, 44-46]. Although results are scarce, it is suspected that extended-spectrum beta-lactamases, such as OXA-48, are found in Lactobacillus isolates like *L. rhamnosus*[12, 46, 47]. Nonetheless, the mechanisms by which Lactobacilli exhibit heteroresistance against cephalosporins remain to be determined. Previous studies using antibiotic susceptibility assays reported that several *Lactobacillus* species are susceptible to tetracyclines and some macrolides[12, 13, 20, 22, 34, 35]. Although most Lactobacilli in this study were susceptible, our work indicates that *L. plantarum* and, *L. reuteri* are heteroresistant to tetracyclines. Our results also demonstrate that *L. gasseri, L. helveticus*, and *L. acidophilus* are heteroresistant to azithromycin. However, all Lactobacilli are resistant to erythromycin, as seen in previous studies[13, 22, 33, 39, 40]. We propose that macrolide-resistance genes, such as the *ermB* resistance gene found against erythromycin, promote Lactobacilli heteroresistance against azithromycin. Nevertheless, the specific heteroresistant mechanisms in Lactobacilli against azithromycin remain to be elucidated.

On the other hand, studies showed that Lactobacilli are resistant to vancomycin, aminoglycosides, and fluoroquinolones[12, 13, 20-22, 32-34, 37, 38]. As expected, most Lactobacilli had a distinct resistant/susceptible phenotype against vancomycin, where half of the Lactobacilli were resistant, and the other half were susceptible. Studies using antibiotic susceptibility assays demonstrated that some Lactobacilli, such as *L. acidophilus, L. plantarum*, and *L. rhamnosus GG*, have varied resistance against aminoglycosides like streptomycin and gentamycin[13, 22]. We confirmed that the varied resistance seen in some Lactobacilli can be categorized as heteroresistant against streptomycin and gentamycin. Our results demonstrated that most Lactobacilli are heteroresistant against fluoroquinolones, especially ciprofloxacin and levofloxacin. However, these Lactobacilli were susceptible to the newer fluoroquinolones, such as delafloxacin. We expect that delafloxacin affects Lactobacilli growth conditions since it lacks a basic group at the C7 position and has chlorine at the C-8 position in its structure[36], rendering it a weak acid that enhances its activity in acidic environments and against Gram-positive bacteria.

Taken together, this work highlights the presence of heteroresistant subpopulations in Lactobacilli, which is widespread among different antibiotic classes. We propose that heteroresistance is a biological feature that *Lactobacillus* species developed for them to live in a complex environment such as the intestines[1, 11, 48]. This study highlights the urgent need to study heteroresistance in commensal microbes.

## Acknowledgments

We want to thank the Emory Antibiotic Resistance Center for their support. This work was supported by startup funds from Emory University (to L.C.-B.) and the following NIH grants: 5R01DK129950-02 (NIDDK; to L.C.-B.), 5R01AI148661-03 (NIAID; to D.W.), U19AI158080 (NIAID; to D.W.) and 1F31AI183823-01 (to D.R.-R.). Additional support was provided by the Emory Antimicrobial Resistance and Therapeutic Discovery Training Program (ARTDTP) under awards 5T32AI106699-08 & 5T32AI106699-09 (to D.R.-R.), and by the Emory Initiative to Maximize Student Development (IMSD) under awards 5R25GM125598-03 & 5R25GM125598-04 (to D.R.-R.). C.B. was supported by the Frances Marx Shillinglaw Women in Science Endowed Fund, the Sherman Fairchild Foundation Summer Stipend Program, the Goizueta Foundation STEM Success Initiative, and the Maria Wornom Rippe ‘64 STEM Research Endowed Fund.

## Author Contributions

D.E.R.-R. designed and performed the experiments, analyzed data, and wrote the paper. C.B. performed experiments and analyzed data. L.C.-B. and D.W. designed the study concept, designed experiments, analyzed experiments, and wrote the paper. All authors reviewed and approved the final submitted manuscript.

## References

1. Donaldson, G.P., S.M. Lee, and S.K. Mazmanian, Gut biogeography of the bacterial microbiota. Nat Rev Microbiol, 2016. 14(1): p. 20–32.

2. Kamada, N., et al., Control of pathogens and pathobionts by the gut microbiota. Nat Immunol, 2013. 14(7): p. 685–90.

3. Latousakis, D., et al., Serine-rich repeat protein adhesins from Lactobacillus reuteri display strain specific glycosylation profiles. Glycobiology, 2019. 29(1): p. 45–58.

4. Sequeira, S., et al., Structural basis for the role of serine-rich repeat proteins from Lactobacillus reuteri in gut microbe-host interactions. Proc Natl Acad Sci U S A, 2018. 115(12): p. E2706–e2715.

5. Thursby, E. and N. Juge, Introduction to the human gut microbiota. Biochem J, 2017. 474(11): p. 1823–1836.

6. Gu, S., et al., Bacterial community mapping of the mouse gastrointestinal tract. PLoS One, 2013. 8(10): p. e74957.

7. MacKenzie, D.A., et al., Strain-specific diversity of mucus-binding proteins in the adhesion and aggregation properties of Lactobacillus reuteri. Microbiology (Reading), 2010.156(Pt 11): p. 3368–3378.

8. Konstantinidis, T., et al., Effects of Antibiotics upon the Gut Microbiome: A Review of the Literature. Biomedicines, 2020. 8(11).

9. Heeney, D.D., M.G. Gareau, and M.L. Marco, Intestinal Lactobacillus in health and disease, a driver or just along for the ride? Curr Opin Biotechnol, 2018. 49: p. 140–147.

10. Salvetti, E. and P.W. O’Toole, The Genomic Basis of Lactobacilli as Health-Promoting Organisms. Microbiol Spectr, 2017. 5(3).

11. Liévin-Le Moal, V. and A.L. Servin, Anti-infective activities of lactobacillus strains in the human intestinal microbiota: from probiotics to gastrointestinal anti-infectious biotherapeutic agents. Clin Microbiol Rev, 2014. 27(2): p. 167–99.

12. Anisimova, E., et al., Alarming Antibiotic Resistance of Lactobacilli Isolated from Probiotic Preparations and Dietary Supplements. Antibiotics (Basel), 2022. 11(11).

13. Campedelli, I., et al., Genus-Wide Assessment of Antibiotic Resistance in Lactobacillus spp. Appl Environ Microbiol, 2019. 85(1).

14. Blaser, M.J., Antibiotic use and its consequences for the normal microbiome. Science, 2016. 352(6285): p. 544–5.

15. Cox, L.M., et al., Altering the intestinal microbiota during a critical developmental window has lasting metabolic consequences. Cell, 2014. 158(4): p. 705–721.

16. Duan, H., et al., Antibiotic-induced gut dysbiosis and barrier disruption and the potential protective strategies. Crit Rev Food Sci Nutr, 2022. 62(6): p. 1427–1452.

17. Panda, S., et al., Short-term effect of antibiotics on human gut microbiota. PLoS One, 2014. 9(4): p. e95476.

18. Shah, T., et al., The Intestinal Microbiota: Impacts of Antibiotics Therapy, Colonization Resistance, and Diseases. Int J Mol Sci, 2021. 22(12).

19. Zhu, S., et al., Assessment of oral ciprofloxacin impaired gut barrier integrity on gut bacteria in mice. Int Immunopharmacol, 2020. 83: p. 106460.

20. Guo, H., et al., Characterization of Antibiotic Resistance Genes from Lactobacillus Isolated from Traditional Dairy Products. J Food Sci, 2017. 82(3): p. 724–730.

21. Rozman, V., et al., Characterization of antimicrobial resistance in lactobacilli and bifidobacteria used as probiotics or starter cultures based on integration of phenotypic and in silico data. Int J Food Microbiol, 2020. 314: p. 108388.

22. Sharma, P., et al., Antibiotic Resistance of Lactobacillus sp. Isolated from Commercial Probiotic Preparations. Journal of Food Safety, 2016. 36(1): p. 38–51.

23. Band, V.I., et al., Antibiotic combinations that exploit heteroresistance to multiple drugs effectively control infection. Nat Microbiol, 2019. 4(10): p. 1627–1635.

24. Band, V.I., et al., Antibiotic failure mediated by a resistant subpopulation in Enterobacter cloacae. Nat Microbiol, 2016. 1(6): p. 16053.

25. Band, V.I. and D.S. Weiss, Heteroresistance to beta-lactam antibiotics may often be a stage in the progression to antibiotic resistance. PLoS Biol, 2021. 19(7): p. e3001346.

26. Dubin, K. and E.G. Pamer, Enterococci and Their Interactions with the Intestinal Microbiome. Microbiol Spectr, 2014. 5(6).

27. Chaguza, C., et al., The population-level impact of Enterococcus faecalis genetics on intestinal colonization and extraintestinal infection. Microbiol Spectr, 2023. 11(6): p. e0020123.

28. Torres, C., et al., Antimicrobial Resistance in Enterococcus spp. of animal origin. Microbiol Spectr, 2018. 6(4).

29. Smith, A.B., et al., Enterococci enhance Clostridioides difficile pathogenesis. Nature, 2022. 611(7937): p. 780–786.

30. Peng, Z., et al., Update on Antimicrobial Resistance in Clostridium difficile: Resistance Mechanisms and Antimicrobial Susceptibility Testing. J Clin Microbiol, 2017. 55(7): p. 1998–2008.

31. Sholeh, M., et al., Antimicrobial resistance in Clostridioides (Clostridium) difficile derived from humans: a systematic review and meta-analysis. Antimicrob Resist Infect Control, 2020. 9(1): p. 158.

32. Klare, I., et al., Antimicrobial susceptibilities of Lactobacillus, Pediococcus and Lactococcus human isolates and cultures intended for probiotic or nutritional use. J Antimicrob Chemother, 2007. 59(5): p. 900–12.

33. Yarahmadi, N., et al., Prevalence of Antibiotic-Resistant Lactobacilli in Sepsis Patients with Long-Term Antibiotic Therapy. Curr Microbiol, 2022. 79(10): p. 318.

34. Goldstein, E.J., K.L. Tyrrell, and D.M. Citron, Lactobacillus species: taxonomic complexity and controversial susceptibilities. Clin Infect Dis, 2015. 60 Suppl 2: p. S98–107.

35. Salminen, M.K., et al., Lactobacillus bacteremia, species identification, and antimicrobial susceptibility of 85 blood isolates. Clin Infect Dis, 2006. 42(5): p. e35–44.

36. Pham, T.D.M., Z.M. Ziora, and M.A.T. Blaskovich, Quinolone antibiotics. Medchemcomm, 2019. 10(10): p. 1719–1739.

37. Zhang, S., et al., d-Alanyl-d-Alanine Ligase as a Broad-Host-Range Counterselection Marker in Vancomycin-Resistant Lactic Acid Bacteria. J Bacteriol, 2018. 200(13).

38. van Pijkeren, J.P. and R.A. Britton, High efficiency recombineering in lactic acid bacteria. Nucleic Acids Res, 2012. 40(10): p. e76.

39. Das, D.J., et al., Critical insights into antibiotic resistance transferability in probiotic Lactobacillus. Nutrition, 2020. 69: p. 110567.

40. Drago, L., et al., Macrolide resistance and in vitro selection of resistance to antibiotics in Lactobacillus isolates. J Microbiol, 2011. 49(4): p. 651–6.

41. Brauner, A., et al., Distinguishing between resistance, tolerance and persistence to antibiotic treatment. Nat Rev Microbiol, 2016. 14(5): p. 320–30.

42. Latousakis, D. and N. Juge, How Sweet Are Our Gut Beneficial Bacteria? A Focus on Protein Glycosylation in Lactobacillus. Int J Mol Sci, 2018. 19(1).

43. Latousakis, D., et al., Serine-rich repeat proteins from gut microbes. Gut Microbes, 2020. 11(1): p. 102–117.

44. Vanichanan, J., et al., Carbapenem-resistant Lactobacillus intra-abdominal infection in a renal transplant recipient with a history of probiotic consumption. Infection, 2016. 44(6): p. 793–796.

45. Ammor, M.S., A.B. Flórez, and B. Mayo, Antibiotic resistance in non-enterococcal lactic acid bacteria and bifidobacteria. Food Microbiol, 2007. 24(6): p. 559–70.

46. Hazırolan, G., et al., Presence of OXA-48 Gene in a Clinical Isolate of Lactobacillus rhamnosus. Foodborne Pathog Dis, 2019. 16(12): p. 840–843.

47. Nouri Gharajalar, S. and M. Firouzamandi, Molecular Detection of Antibiotic Resistance Determinants in Lactobacillus Bacteria Isolated from Human Dental Plaques. Journal of Medical Microbiology and Infectious Diseases, 2017. 5(3): p. 51–55.

48. Kim, S., A. Covington, and E.G. Pamer, The intestinal microbiota: Antibiotics, colonization resistance, and enteric pathogens. Immunol Rev, 2017. 279(1): p. 90–105.

