## Supplementary materials for "Widespread heteroresistance to antibiotics in Lactobacillus species"

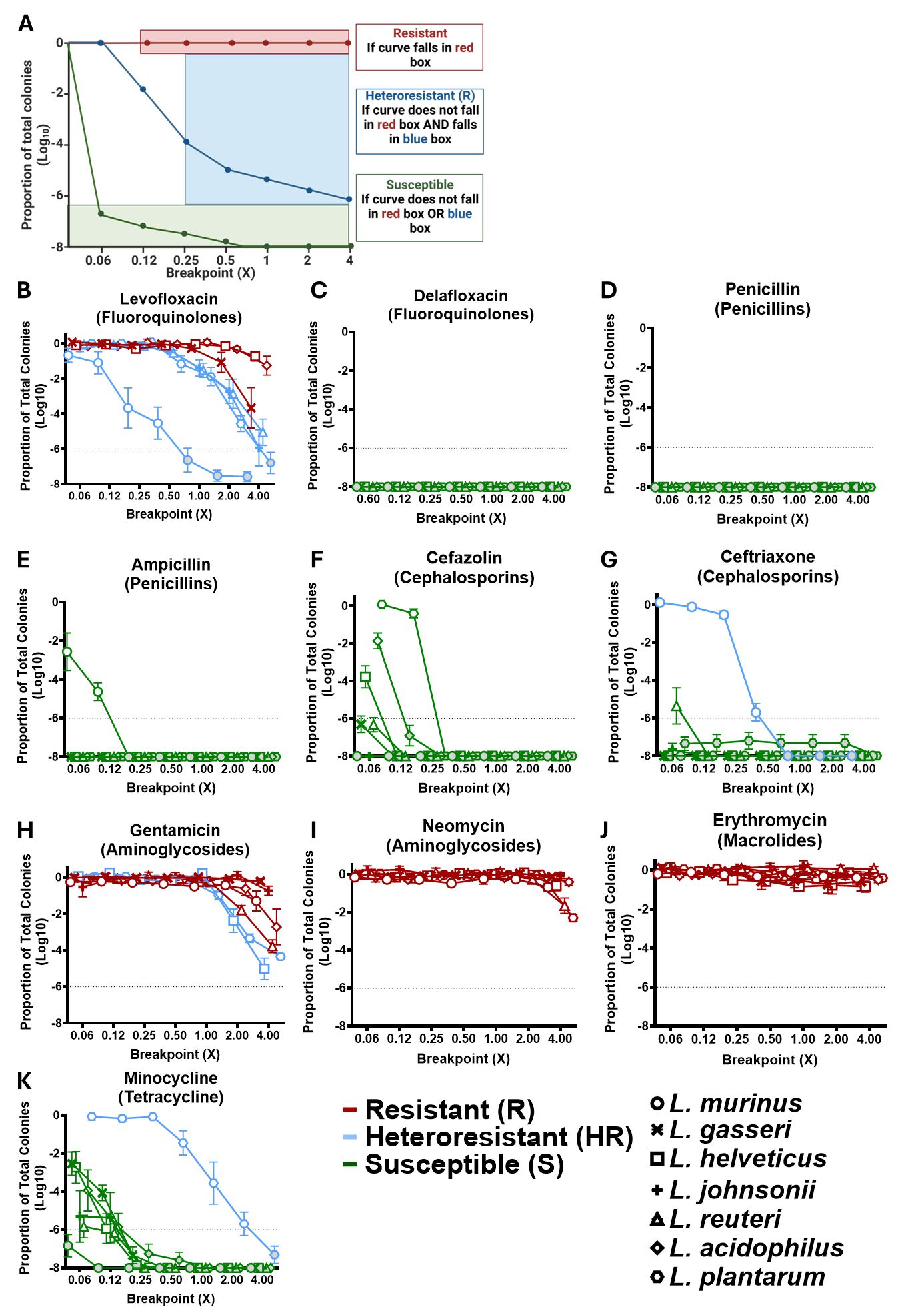

**Figure S1: Lactobacilli have varied resistance to different antibiotic classes. A)** Representative population analysis profiles (PAP) graph for **resistant (R)**, **heteroresistant (HR)**, or **susceptible (S)** isolates, as well as graph sectors from which designations were made. Population analysis profile assays (PAPs) of seven Lactobacillus isolates plated on **B-C)** Fluoroquinolones (**B**: Levofloxacin; MIC: 8µg/ml, **C**: Delafloxacin; MIC: 4µg/ml), **D-E)** Penicillins (**D**: Penicillin; MIC: 16µg/ml, **E**: Ampicillin; MIC: 16µg/ml), **F-G)** Cephalosporins (**F**: Cefazolin; MIC: 64µg/ml, **G**: Ceftriaxone; MIC: 64µg/ml), **H-I)** Aminoglycosides (**H**: Gentamicin; MIC: 16µg/ml, **I**: Neomycin; MIC: 16µg/ml), **J)** Macrolides (Erythromycin; MIC: 8µg/ml), **K)** Tetracyclines (Minocycline; MIC: 16µg/ml) in aerobic conditions. **Gray-filled** dots indicate samples below the level of detection. Symbols represent bacterial species. Graphs are representative of 3 independent experiments (n=3).

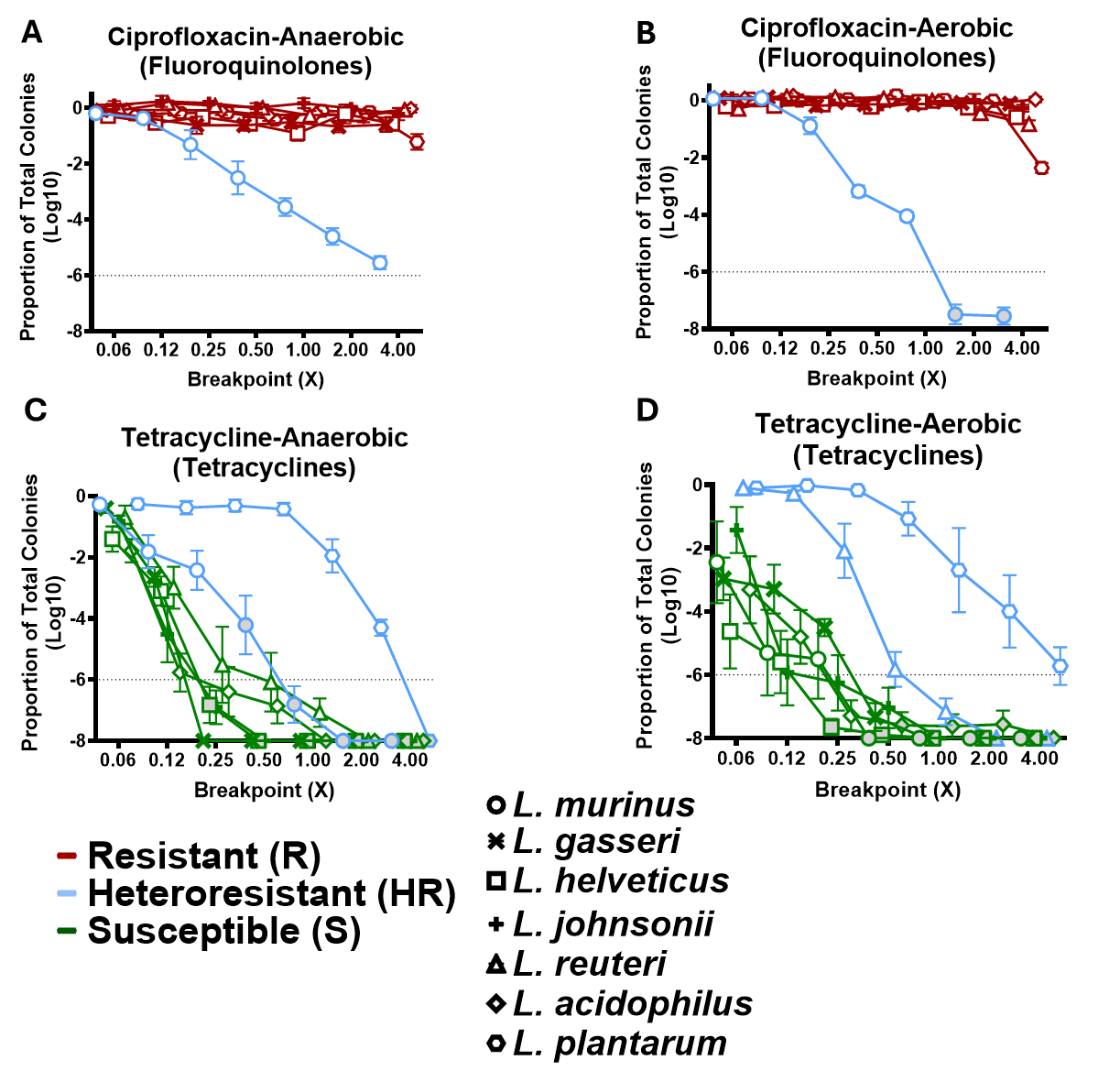

**Figure S2: Lactobacilli have varied resistance to different antibiotic classes in anaerobic conditions.** Population analysis profiles (PAPs) of Lactobacillus isolates plated on **A-B)** Ciprofloxacin (MIC: 4µg/ml), **C-D)** Tetracycline (MIC: 16µg/ml) in **A & C)** anaerobic and **B & D)** aerobic conditions. **Gray-filled** dots indicate samples below the level of detection. Symbols represent bacterial species. Graphs are representative of 3 independent experiments (n=3).

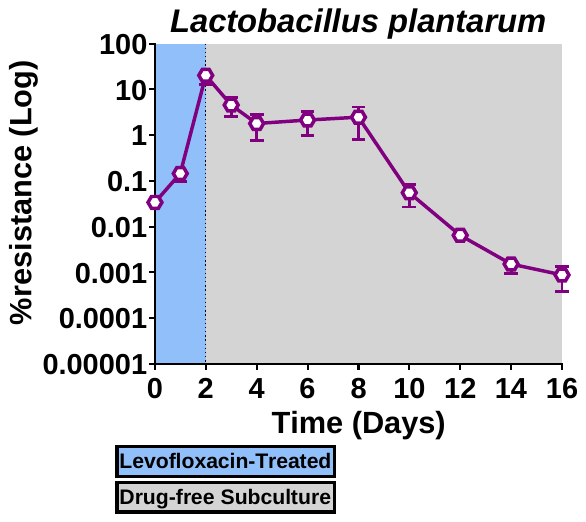

**Figure S3: *Lactobacillus plantarum* exhibits heteroresistance to levofloxacin.** *L. plantarum* was subcultured in MRS with levofloxacin (MIC: 8µg/ml) for 48 hours followed by subcultures in levoxacin-free MRS media for 14 days. Data represents the proportion of levofloxacin resistant bacteria in the culture each day. Symbols represent the mean of triplicates and is representative of three independent experiments (n=9).

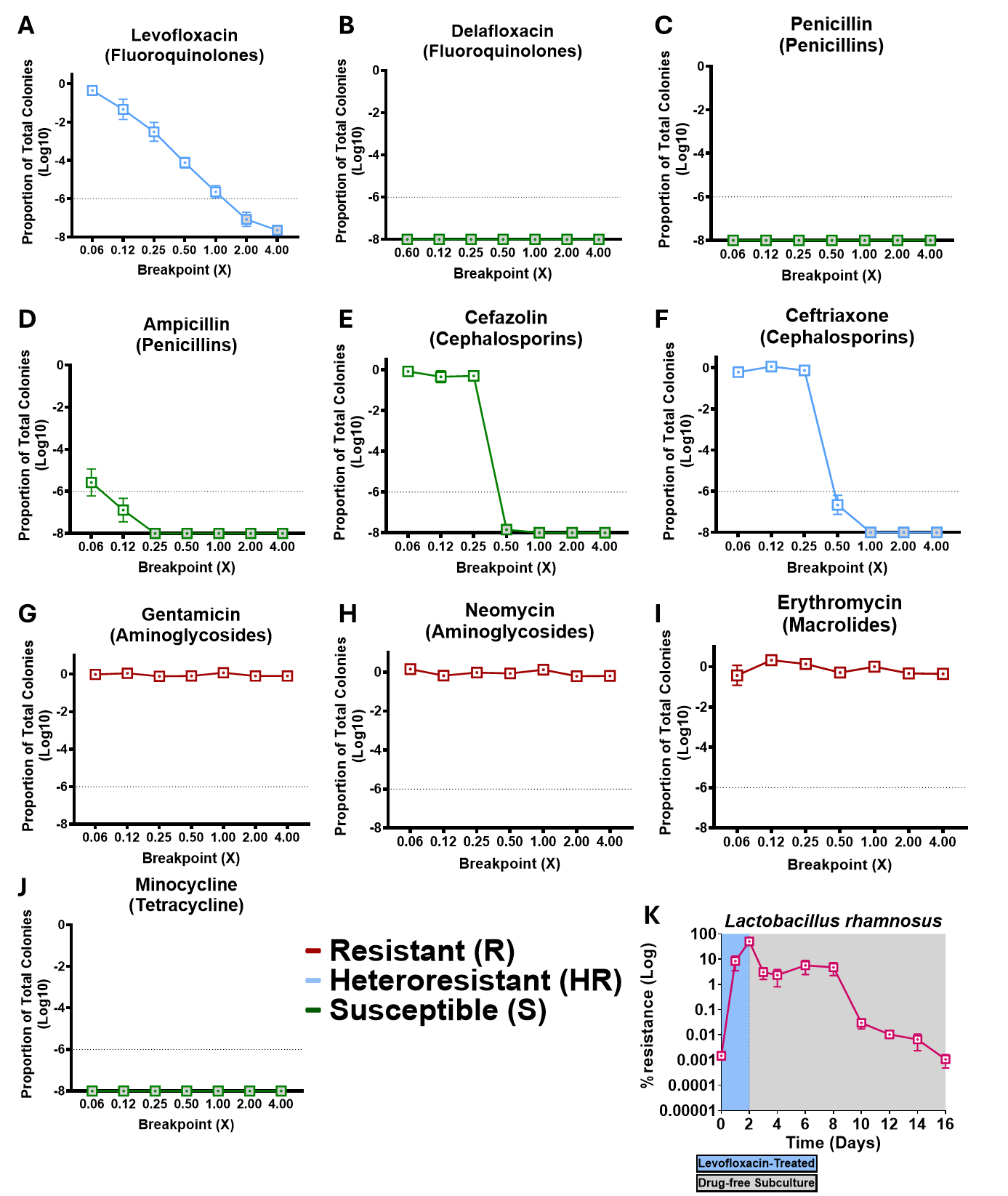
experiments (n=9).

**Figure S4*: Lactobacillus rhamnosus* has varied resistance to different antibiotics.** Population analysis profiles assay (PAPs) of *L. rhamnosus* plated on **A-B**) Fluoroquinolones (**A**: Levofloxacin; MIC: 8µg/ml, **B**: Delafloxacin; MIC: 4µg/ml), **C-D**) Penicillins (**C**: Penicillin; MIC: 16µg/ml, **D**: Ampicillin; MIC: 16µg/ml), **E-F**) Cephalosporins (**E**: Cefazolin; MIC: 64µg/ml, **F**: Ceftriaxone; MIC: 64µg/ml), **G-H**) Aminoglycosides (**G**: Gentamicin; MIC: 16µg/ml, **H**: Neomycin; MIC: 16µg/ml), **I**) Macrolides (Erythromycin; MIC: 8µg/ml), **J**) Tetracyclines (Minocycline; MIC: 16µg/ml) in aerobic conditions. Gray-filled dots indicate samples below the level of detection. Symbols represent bacterial species. Graphs are representative of 3 independent experiments (n=3).**K)** *L. rhamnosus* was subcultured in MRS with levofloxacin (MIC: 8µg/ml) for 48 hours followed by subcultures in levofloxacin-free MRS media for 14 days. Data represents the proportion of levofloxacin resistant bacteria in the culture each day. Symbols represent the mean of triplicates and is representative of three independent experiments (n=9).

**Table S1 Antibiotics table**

| **Fluoroquinolones** | | | |
| --- | --- | --- | --- |
| **Name** | **Company** | **Catalog Number** | **Storage Temperature (°C)** |
| Ciprofloxacin HCl | Alfa Aesar | J61317-06 | -20 |
| Levofloxacin | Sigma | 28266-10G-F | 4 |
| Delafloxacin meglumine | Combi Blocks | HG-0484 | -20 |

| **Penicillins** | | | |
| --- | --- | --- | --- |
| **Name** | **Company** | **Catalog Number** | **Storage Temperature (°C)** |
| Ampicillin Sodium Salt | Research Product International | A40040-25.0 | 4 |
| Amoxicillin Sodium Salt | Alfa Aesar | J66675 | 4 |
| Penicillin G Sodium Salt | Alfa Aesar | J63032 | 4 |

| **Macrolides** | | | |
| --- | --- | --- | --- |
| **Name** | **Company** | **Catalog Number** | **Storage Temperature (°C)** |
| Azithromycin Dihydrate | Astatech | 44144 | 4 |
| Erythromycin | Sigma | E5389-1G | 20 |

| **Cephalosporins** | | | |
| --- | --- | --- | --- |
| **Name** | **Company** | **Catalog Number** | **Storage Temperature (°C)** |
| Cefepime Hydrochloride Monohydrate | Alfa Aesar | J66237 | 4 |
| Cefazolin Sodium Salt | TCI | C2242 | 4 |
| Ceftriaxone Disodium Salt | TCI | C2226 | 4 |

| **Aminoglycosides** | | | |
| --- | --- | --- | --- |
| **Name** | **Company** | **Catalog Number** | **Storage Temperature (°C)** |
| Streptomycin Sulfate | Gibco | 11860038 | 4 |
| Neomycin Sulfate | CalbioChem | 4801 | 20 |
| Gentamycin Sulfate | MP Biochemicals | 190057 | 4 |

| **Tetracyclines** | | | |
| --- | --- | --- | --- |
| **Name** | **Company** | **Catalog Number** | **Storage Temperature (°C)** |
| Tetracycline Hydrochloride | Gibco | A39246 | -20 |
| Minocycline Hydrochloride | Alfa Aesar | J66429 | 4 |

| **Glycopeptides** | | | |
| --- | --- | --- | --- |
| **Name** | **Company** | **Catalog Number** | **Storage Temperature (°C)** |
| Vancomycin hydrochloride | Alfa Aesar | J62790 | 4 |
